# Chaotic dynamics in spatially distributed neuronal networks generate population-wide shared variability

**DOI:** 10.1101/2022.04.19.488846

**Authors:** Noga Mosheiff, Bard Ermentrout, Chengcheng Huang

## Abstract

Neural activity in the cortex is highly variable in response to repeated stimuli. Population recordings across the cortex demonstrate that the variability of neuronal responses is shared among large groups of neurons and concentrates in a low dimensional space. However, the source of the populationwide shared variability is unknown. In this work, we analyzed the dynamical regimes of spatially distributed networks of excitatory and inhibitory neurons. We found chaotic spatiotemporal dynamics in networks with similar excitatory and inhibitory projection widths, an anatomical feature of the cortex. The chaotic solutions contain broadband frequency power in rate variability and have distance-dependent and low-dimensional correlations, in agreement with experimental findings. In addition, rate chaos can be induced by globally correlated noisy inputs. These results suggest that spatiotemporal chaos in cortical networks can explain the shared variability observed in neuronal population responses.

## Introduction

A defining feature of cortical neural responses is that they are highly variable. The variability is reflected at multiple scales of neural recordings, from irregular inter-spike intervals in spike trains of individual neurons (Kara et al., 2000; Shadlen and Newsome, 1998) to spatiotemporal patterns in mesoscopic neural activity measured with voltage sensitive dye imaging (Arieli et al., 1996) and local field potentials (Muller et al., 2014), to whole brain signals such as the electroencephalography (Waschke et al., 2019). Changes in neural variability reflect fluctuations in the brain state and are closely related to behavioral performance (Doiron et al., 2016; McGinley et al., 2015; Ni et al., 2018; Waschke et al., 2019). Therefore, understanding the circuit mechanisms that generate neural variability is critical for elucidating the neural basis of behavior.

Previous models have proposed that chaotic neural dynamics can be a substantial local source of neural variability in cortical circuits (Rajan et al., 2010; Sompolinsky et al., 1988; Van Vreeswijk and Sompolinsky, 1996). Variable neural responses can be intrinsically generated through strong interactions between the excitatory and inhibitory neurons. Intriguingly, neuronal networks with chaotic dynamics have been shown to demonstrate high computational capabilities because of their rich reservoir of internal dynamics that can be utilized for complex computations and efficient training (Bertschinger and Natschläger, 2004; Jaeger and Haas, 2004; Laje and Buonomano, 2013; Sussillo and Abbott, 2009; Toyoizumi and Abbott, 2011).

However, previous models with unstructured random connectivity produce chaotic response in individual neurons that is uncorrelated with other neurons in the network. In contrast, numerous datasets of cortical recordings have revealed that cortical neurons are on average positively correlated (Cohen and Kohn, 2011). The correlation between a pair of neurons depends on many factors, such as the cortical distance between them and their tuning similarity (Lin et al., 2015; Schulz et al., 2015; Smith and Sommer, 2013). Moreover, the variability shared among a neuron population has been found to be low dimensional, meaning that the variations in population response patterns can often be explained by just a few independent latent variables (Huang et al., 2019; Ruff et al., 2020; Schölvinck et al., 2015; Semedo et al., 2019; Williamson et al., 2016). Therefore, networks with unstructured random connectivity are not able to capture the shared variability in neural population responses.

A main determinant of the connection probability between a pair of neurons in cortex is the physical distance between them (Levy and Reyes, 2012; Mariño et al., 2005a; Oswald et al., 2009; Oswald and Reyes, 2011; Rossi et al., 2020). Nearby neurons are more likely to be connected, whereas neurons that are far apart are less likely to be connected. Recently, several studies of spatially distributed neuronal networks have suggested that spatiotemporal patterns of neural activity can explain many features of variability in neural population responses (Huang et al., 2019; Keane and Gong, 2015; Pyle and Rosenbaum, 2017; Shi et al., 2022). For example, our past work has shown that spiking neuron networks with irregular wave dynamics generate on average positive correlations and low-dimensional population-wide shared variability, consistent with cortical recordings (Huang et al., 2019).

Here we systematically analyze the dynamical regimes of spatially distributed firing rate networks. We find a parameter region where networks exhibit irregular chaotic dynamics. The chaotic solutions have several features of response variability that are consistent with experimental findings in cortex, such as broadband frequency power in single neuron responses, distance-dependent correlations and low-dimensionality of population responses. Interestingly, chaos occurs in networks where the excitatory and inhibitory neurons have similar projection widths, an anatomical feature found in the cortex (Levy and Reyes, 2012; Mariño et al., 2005a; Rossi et al., 2020). Further, we find that correlated noisy inputs induce chaos, which can explain the prevalence of large-scale shared variability observed in cortex. Our work identifies a new dynamical regime of spatiotemporal chaos in neuronal networks that can account for rich response patterns in neural population activity.

## Results

We study a spatially distributed network model that describes the firing rate dynamics of the excitatory (*r*_*e*_) and inhibitory (*r*_*i*_) neurons (Eqs. 1-2). Neurons are organized on a two-dimensional sheet (*x* ∈ [0, 1] × [0, 1]) with periodic boundary conditions (Fig. 1A). The equations that describe the dynamics of the firing rates are

**Figure 1:**
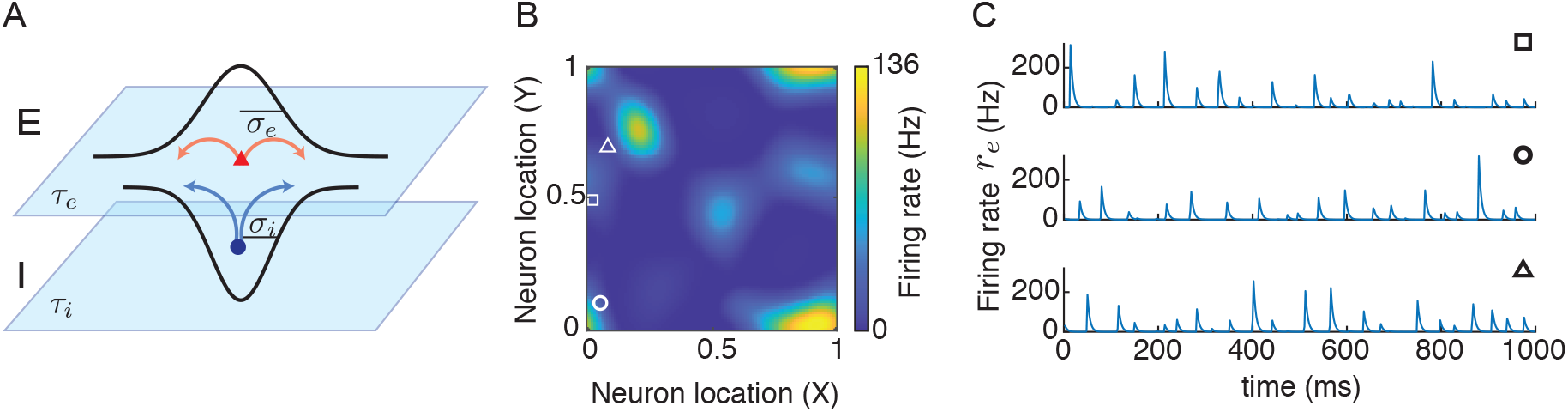
Irregular spatiotemporal dynamics in spatially distributed networks. **A**. Model schematic of recurrently coupled excitatory (E) and inhibitory (I) neurons. Neurons from each population are equally spaced on a two dimensional sheet, [0, 1] × [0, 1], with distance-dependent connectivity weights. **B**. A snapshot of the firing rates of excitatory neurons. **C**. Three examples of the time courses of firing rates of neurons at different locations. The square, circle and triangle in panel **B** denote the spatial location of each neuron.

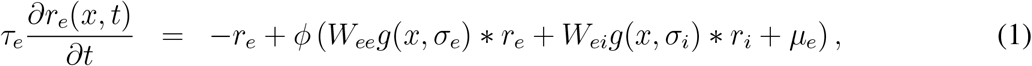

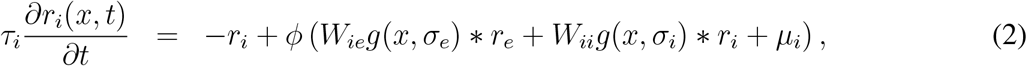

where *τ*_*e*_ (*τ*_*i*_) is the time constant of the excitatory (inhibitory) population, * denotes convolution in space, and *ϕ*(*x*) = max(*x*, 0)^2^ is the input-output transfer function of each neuron. The connection strength from a neuron in population *β* to a neuron in population *α* decays with distance as a Gaussian function, *g*(*x, σ*_*β*_), with projection width *σ*_*β*_ (*α, β* = *e, i*). The average connection strength from population *β* to population *α* is *W*_*αβ*_. The external input to each population is a static and spatially homogeneous current, *µ*_*α*_ (*α* = *e, i*).

The spatial networks generate rich spatiotemporal patterns. In particular, we find that the network exhibits irregular patterns in both space and time for certain parameters (Fig. 1B-C). These networks show spatially localized and transient activity patterns that sometimes propagate across the network (Fig. 1B). Individual neurons show epochs of brief firing with varying magnitudes and time intervals in between (Fig. 1C). This type of network dynamics result in large variability that is shared among neurons in the network. In order to better understand the behavior of the model and the mechanism for generating irregular firing patterns, we systematically analyze the different dynamical regimes of the spatially distributed networks. We focus our analysis on varying the temporal (*τ*_*i*_) and spatial (*σ*_*i*_) scales of the inhibitory neurons, while fixing those of the excitatory neurons (*τ*_*e*_ = 5 ms and *σ*_*e*_ = 0.1).

### A reduced two-unit model with no spatial coupling

We first consider networks with no spatial coupling, meaning that the spatial coupling function, *g*, is constant over distance. In this case, the network is reduced to a two-unit system, where the firing rate, *r*_*α*_(*x, t*) = *r*_*α*_(*t*), *α* = *e, i*, is independent of the neural location *x*.

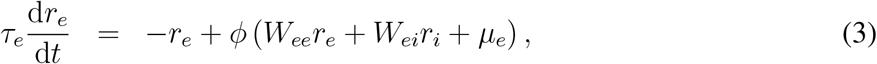

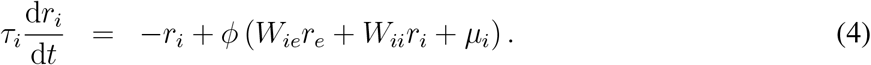

Note that a solution to the two-unit system corresponds to a spatially uniform solution to the spatially distributed networks (Eqs. 1-2).

The reduced network (Eqs. 3-4) has a stable fixed point solution for a small *τ*_*i*_ (Fig. 2A, gray solid line). As *τ*_*i*_ increases, the fixed point solution becomes unstable through a Hopf bifurcation (*τ*_*i*_ = *τ*_HB_, Fig. 2A, gray arrow), and the system admits a stable periodic solution (limit cycle; Fig. 2A, blue solid curve). Over the interval of *τ*_*i*_ ∈ [7.16, 7.78] ms both the fixed point and the limit cycle solutions are stable (between the blue and gray arrows in Fig. 2A). We next analyze the stability of the fixed point and the limit cycle solutions in the spatially distributed networks (Eqs. 1-2).

**Figure 2:**
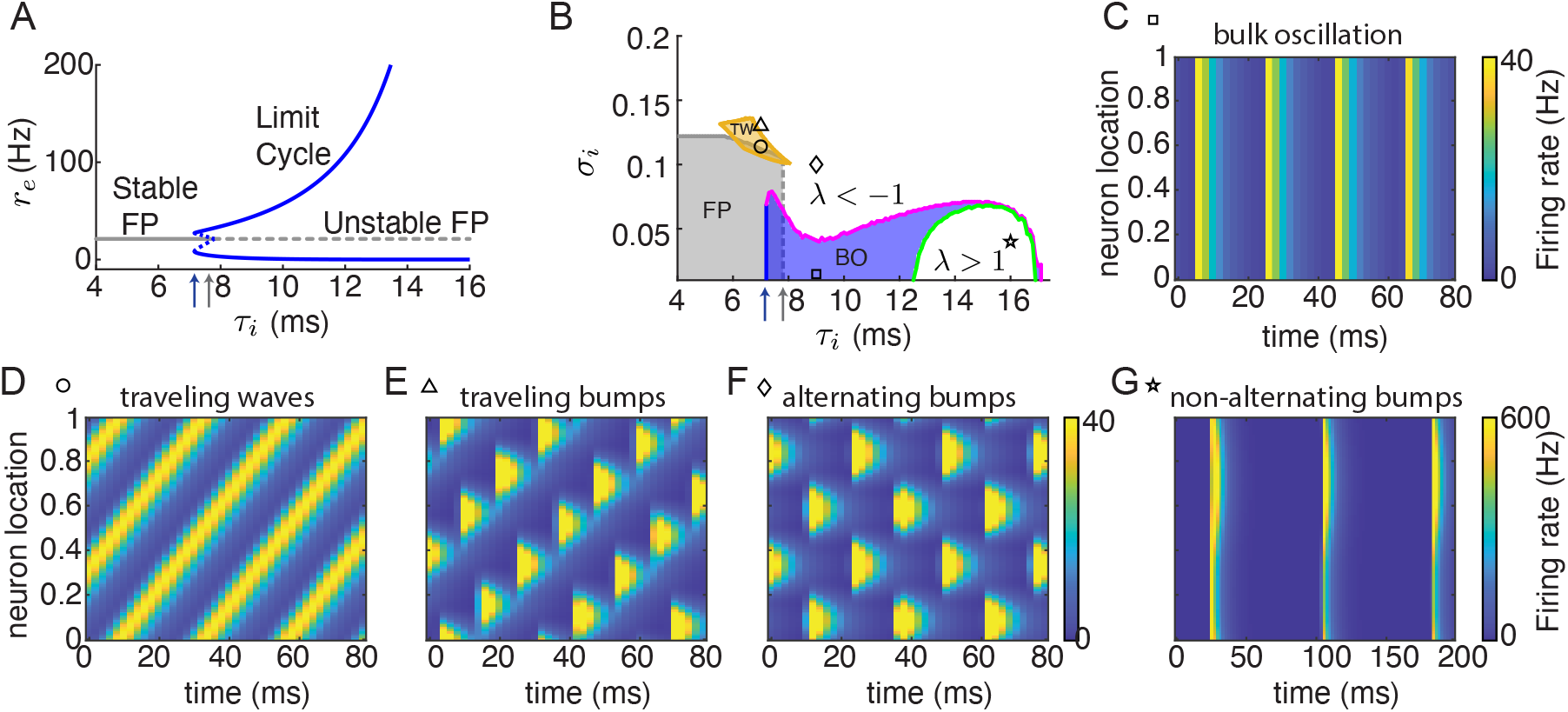
Dynamical regimes of networks with one-dimensional spatial coupling. **A**. Bifurcation diagram of the reduced two-unit model (Eq. 3-4) as *τ*_*i*_ varies. Gray line, fixed point; blue curves, the maximal and minimal firing rates of a limit cycle; solid, stable; dashed, unstable. The blue arrow indicates the lowest *τ*_*i*_ with a limit cycle solution. The gray arrow indicates the Hopf bifurcation (the largest *τ*_*i*_ with a stable fixed point solution). **B**. Phase diagram of networks with one dimensional spatial coupling. Gray, stable fixed point; blue, stable bulk oscillation; orange, stable traveling waves. The fixed point solution loses stability at either the zero wave number (gray dashed line) or a nonzero wave number (gray solid line). The bulk oscillation loses stability with either an eigenvalue becoming larger than 1 (green curve) or an eigenvalue becoming less than −1 (magenta curve). Blue and gray arrows point to the same values of *τ*_*i*_ as in **A. C-G** Examples of the firing rate dynamics of the excitatory neurons in networks with different *τ*_*i*_ and *σ*_*i*_ (the square, circle, triangle, diamond and star symbols indicate the parameters in panel **B**). **C**, Bulk oscillation; **D**, traveling waves; **E**, traveling bumps; **F**, alternating bumps; **G**, non-alternating bumps. The temporal and spatial scales of the excitatory neurons are *τ*_*e*_ = 5 ms and *σ*_*e*_ = 0.1, respectively.

### Pattern formation in one-dimensional networks

In this section we analyze the stability of spatially uniform solutions and how a loss of stability leads to periodic spatiotemporal patterns. We first consider networks with one-dimensional spatial coupling, where neurons are distributed over a line interval, [0, 1], with periodic boundary conditions (namely, a ring). The stability analysis below is also applicable to two-dimensional networks.

We first analyze the stability of the fixed point solution in spatially distributed networks, which is a static and spatially uniform solution. We linearize around the fixed point in the spatial frequency domain using Fourier transform and obtain eigenvalues for each wave number (spatial frequency) (see Methods; Ermentrout and Cowan (1979); Rosenbaum and Doiron (2014)). The fixed point solution is stable when all eigenvalues are negative (stable region is shown in gray in Fig. 2B). When *σ*_*i*_ *< σ*_*e*_, the network loses stability at zero wave number as *τ*_*i*_ increases, which is the same condition as the Hopf bifurcation in the two-unit model (Fig. 2B, gray arrow, gray dashed line). For small *σ*_*i*_ and *τ*_*i*_ *> τ*_HB_, the network exhibits spatially uniform and temporally periodic solutions (bulk oscillation; Fig. 2C), which corresponds to the limit cycle solution in the two-unit model (Eq. 3-4). When *σ*_*i*_ *> σ*_*e*_, the network loses stability at a nonzero wave number (Fig. 2B, gray solid line), suggesting pattern formation of spatially and/or temporally periodic solutions.

Around the boundary where the fixed point solution loses stability at a nonzero wave number, we find traveling wave solutions (Fig. 2B orange region, Fig. 2D). Using a continuation numerical method (Avitabile, 2016), we show that stable traveling wave solutions exist in a small parameter region (Fig. 2B, orange) that partially overlaps with the region of stable fixed point. Closely beyond this region with a larger *σ*_*i*_, the traveling waves lose stability and the networks generate traveling bump solutions (Fig. 2E).

We next compute the stability of the bulk oscillation solution. Similar to the stability analysis of the fixed point solution, we linearize around the bulk oscillation solution and perturb the system at different wave number (see Methods). The dynamics of the perturbation then follow a linear system of differential equations with periodic coefficients (Eq. 8). The stability of the bulk oscillation solution depends on the eigenvalues (*λ*) of the Monodromy matrix, *M*, of the linear system, which describes the change of the perturbation after one period of the bulk oscillation solution (Eq. 9; Ali et al. (2016); Teschl (2012)). The bulk solution is unstable if there is an eigenvalue of *M* (*k*) with magnitude larger than 1 for any wave number *k*. We find that with small *σ*_*i*_ and an intermediate range of *τ*_*i*_, the bulk oscillation is stable (Fig. 2B, blue region; Fig. 2C). As *σ*_*i*_ increases, the bulk oscillation loses stability with a real eigenvalue less than −1 for perturbations at a nonzero wave number, indicating a period-doubling bifurcation (Fig. 2B, magenta curve). In the parameter region beyond this stability boundary, the network activity shows spatial patterns that alternate over time (Fig. 2F). As *τ*_*i*_ increases, the bulk oscillation loses stability with a real eigenvalue larger than 1 (Fig. 2B, green curve). In the region under this stability boundary (Fig. 2B, green curve), the network activity exhibits spatial patterns that repeats at each cycle (Fig. 2G).

### Chaotic dynamics in two dimensional networks

We next analyze the full networks with two dimensional spatial coupling (Fig. 1A). The stability analysis of the fixed point and the bulk oscillation that we outlined above for the one-dimensional networks is also applicable to networks of higher dimensions. The two-dimensional networks have almost identical stability boundaries for the fixed point and the bulk oscillation solutions as those in the one-dimensional networks (Fig. 2B and Supplemental Fig. S1). In the region above the period-doubling bifurcation curve of the bulk oscillation solution (Fig. 2B, region with *λ <* −1), the two-dimensional networks have solutions similar to those in the one-dimensional networks, such as traveling waves (Fig. 3A) and alternating bumps solutions (Fig. 3B). In addition, we find other spatiotemporal patterns in the two-dimensional networks, such as spatially periodic stripe patterns that alternate in phase over time (Fig. 3C).

**Figure 3:**
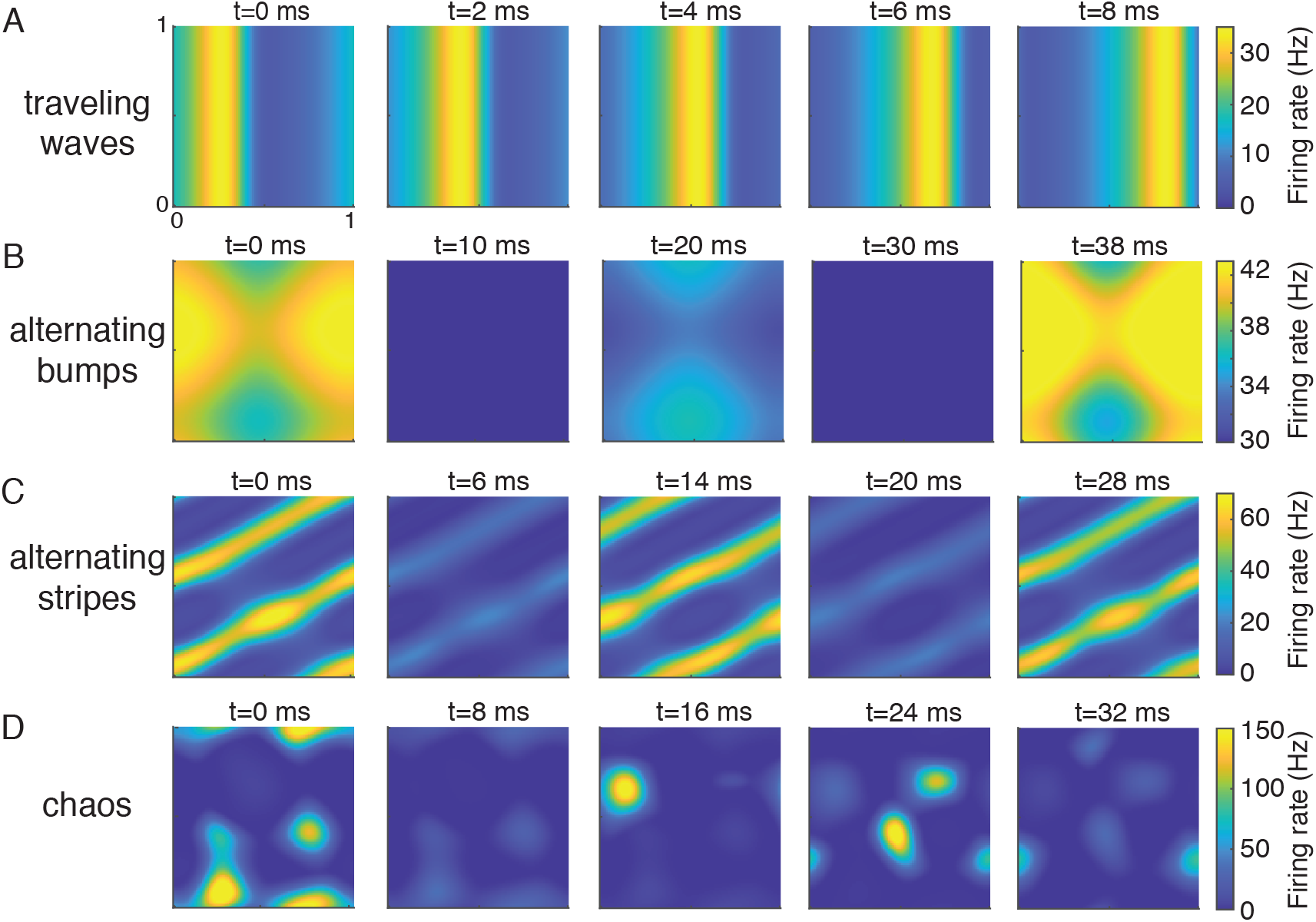
Examples of solutions in networks with two-dimensional spatial coupling. **A-D**, Snapshots of the firing rates of the excitatory population at five time frames. **A**, traveling waves solution (*τ*_*i*_ = 8, *σ*_*i*_ = 0.1). **B**, alternating bumps solution (*τ*_*i*_ = 9, *σ*_*i*_ = 0.06). **C**, alternating stripes solution (*τ*_*i*_ = 9, *σ*_*i*_ = 0.1). **D**, chaotic solution (*τ*_*i*_ = 12.8, *σ*_*i*_ = 0.096). The axes are the neuron location on the two-dimensional neuronal sheet.

In particular, we find network solutions that are irregular in both space and time (Fig. 3D, Fig. 1B,C). In these networks, neurons exhibit random like activity with large variability even though the network is deterministic. We verify that such irregular solutions are chaotic, meaning that a small perturbation leads to rapid divergence in network activity, by computing the maximal Lyapunov exponent (MLE) numerically (see Methods; Wolf et al. (1985)). A positive MLE means that the solution is chaotic, a negative MLE indicates a stable fixed point, and MLE=0 indicates that the solution is periodic or quasi-periodic. We compute the MLE of network solutions over the parameter space of *σ*_*i*_ and *τ*_*i*_. We find chaotic solutions in a parameter region where the spatial scales of the excitatory and inhibitory projections are similar (*σ*_*i*_ ≈ *σ*_*e*_ = 0.1) and the time constant of the inhibitory neurons is large (*τ*_*i*_ *> τ*_*e*_ = 5 ms) (Fig. 4A, yellow). Interestingly, anatomical measurements from sensory cortices find that excitatory and inhibitory neurons project on similar spatial scales (Levy and Reyes, 2012; Mariño et al., 2005b; Rossi et al., 2020). In addition, the decay kinetics of inhibitory synaptic currents are also slower than the excitatory synaptic currents in physiology (Geiger et al., 1997; Oswald and Reyes, 2011; Xiang et al., 1998). These results suggest that the network parameter region of chaotic dynamics is consistent with the anatomy and physiology of the cortex. In contrast, networks with one-dimensional spatial coupling do not generate chaos in this parameter region, suggesting that the two-dimensional spatial structure is important for generating chaos (Supplementary Fig. S2).

**Figure 4:**
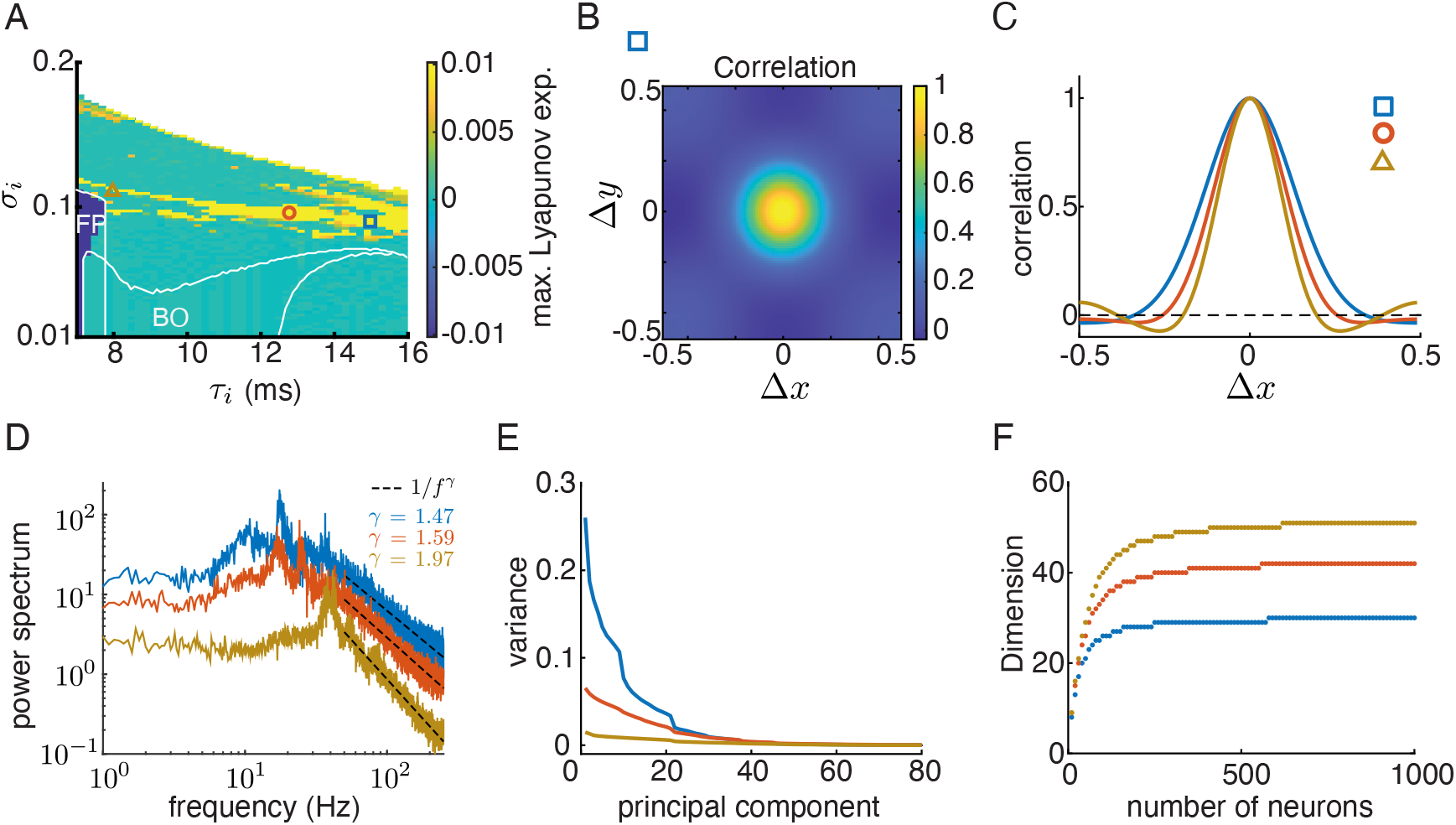
Population statistics of the chaotic solutions. **A**. The maximal Lyapunov exponent (MLE) as a function of the projection width (*σ*_*i*_) and the time constant (*τ*_*i*_) of the inhibitory population for the two-dimensional networks. The white curves are the stability borders of the parameter regions of stable fixed point (FP) and bulk oscillation (BO) solutions (same regions as in Fig. 2B and Supplemental Fig. S1). **B**. The correlation between neuron activity as a function of distance in *x* and *y* directions (Eq. 11) in a network labeled as a blue square in panel A. **C**. The correlation function along the *x* direction for three networks in the chaotic regime (Δ*y* = 0). The network parameters are denoted in panel A with corresponding symbols. **D**. Population averaged power spectrum of single-neuron rate activity represented in log-log scale (Eq. 12). Dashed lines represent 1*/f*^*γ*^ scaling. **E**. The variance of the dominant principal components of population rate activity of 1000 randomly sampled neurons from the networks, averaged over 100 sampling. **F**. The dimension of population activity saturates as the number of sampled neurons increases. The dimension is defined as the number of dominant principal components that account for 95% of the total variance.

### Population statistics of the chaotic solutions

Networks with chaotic dynamics intrinsically generate large variability in population activity. We next measure statistics of the population activity of networks in the chaotic regime and compare with cortical recordings. First, we compute the spatial correlations of population activity along *x* and *y* directions (Fig. 4B-C, see Methods). The spatial correlations of the chaotic dynamics are roughly isotropic, and decrease with the distance between neurons (Fig. 4B). The decrease of correlation with cortical distance has been widely reported in population recordings across the cortex (Nauhaus et al., 2009; Pinto and Dan, 2015; Rothschild et al., 2010; Smith et al., 2018; Smith and Sommer, 2013; Yu et al., 2019). Further, we found that the spatial decay rate of correlation depends on network parameters. Networks with larger *τ*_*i*_ have a broader range of correlation which remains positive over long distance (Fig. 4C).

Second, we measure the power spectrum of the temporal variability of each neuron unit. We find that neurons in chaotic networks have broadband frequency power with peaks at around 20-40 Hz (Fig. 4D). Beyond about 50 Hz, the power decays with frequency following a power law scaling (1*/f*^*γ*^) with exponent (*γ*) between 1.5 and 2 (Fig. 4D dashed). This means that the neurons’ rate activities are not periodic as those in regular solutions, but exhibit considerable variability over a broad frequency range. This type of broadband frequency power has also been found in many neural recordings (Henrie and Shapley, 2005; Okun et al., 2019; Ray and Maunsell, 2010).

Lastly, we examine the dimensionality of the chaotic solutions. We perform principal component analysis of the rate activity of 1000 randomly sampled neurons from the networks. The variance of each principal component is the eigenvalue of the covariance matrix of population rate over time, sorted from large to small. We find that the variability of the chaotic population rate concentrates in the first few tens of principal components (Fig. 4E). We define the dimension of the population activity as the number of principal components that explains 95% of the variance. The dimension of chaotic solutions increases with the number of sampled neurons and saturates below 50 (Fig. 4F), which is much lower than the number of neurons in the network (*N* = 10^4^). The low dimensional structure is a defining feature of cortical neural variability that has been recently demonstrated in multiple population recordings (Huang et al., 2019; Lin et al., 2015; Ruff et al., 2020; Schölvinck et al., 2015; Williamson et al., 2016). In contrast, the chaotic rate dynamics in disordered random networks have dimensions that increase linearly with network size (Engelken et al., 2020).

### Correlated input noise expands the chaotic regime

We have, thus far, analyzed the behavior of fully deterministic networks. We now consider how input noise changes network dynamics. The input to neuron *k* from population *α* is

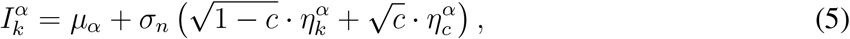

where *σ*_*n*_ is the standard deviation and *c* ∈ [0, 1] is the correlation coefficient of the input noise. The input noise to each neuron consists of two components: 1) an independent noise component, 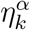, which is private to each neuron *k*, and 2) a correlated noise component, 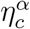, which is common for all neurons in population *α* ∈ {*e, i*} (Fig. 5A; De La Rocha et al. (2007); Kanashiro et al. (2017)). Both noise components are modeled as Ornstein–Uhlenbeck processes with time constant *τ*_*n*_ = 5 ms (see Methods, Eq. 13). The amplitude of the independent noise component is 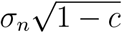 and the amplitude of the correlated component is 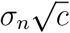.

**Figure 5:**
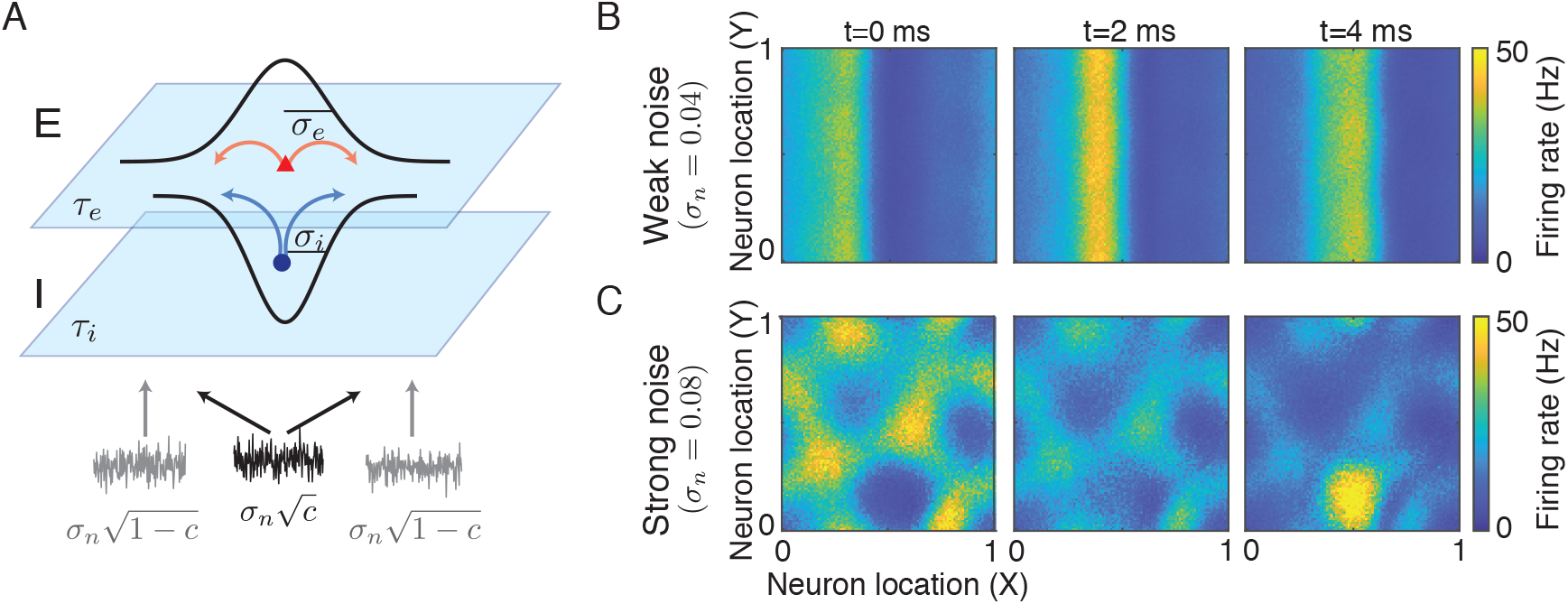
Two-dimensional networks with input noise. **A**. The input noise to each neuron consists of a correlated noise component (black) that is common for all neurons from the same population, and an independent noise component that is private to each neuron (gray). **B-C**. Snapshots of the firing rates of the excitatory population at three time frames for a network that generates traveling waves when there is no input noise (*τ*_*i*_ = 8, *σ*_*i*_ = 0.1, same parameters as in Fig. 3A). The correlation of the input noise is *c* = 0.5, and the amplitude is *σ*_*n*_ = 0.04 (**B**) and *σ*_*n*_ = 0.08 (**C**).

When the input noise is weak (small *σ*_*n*_), the regular solutions can roughly maintain their spatiotemporal patterns (Fig. 5B). However, when the input noise is strong (large *σ*_*n*_), the patterns might be distorted or completely destroyed. For example, a network that produces traveling waves without input noise (Fig. 3A) can still generate a noisy traveling wave pattern with weak noise (Fig. 5B), but exhibits irregular patterns when input noise is strong (Fig. 5C).

The irregular spatiotemporal patterns in networks with strong noise are similar to the chaotic solutions in deterministic networks (Fig. 3A). We next compute the MLE of networks with frozen input noise in the parameter space of *σ*_*i*_ and *τ*_*i*_ (see Methods). We find that input noise can induce chaos in the spatially distributed networks. With correlated input noise, the parameter region of chaotic solutions expands as the amplitude of noise increases (compare yellow regions in Fig. 4A and Fig. 6A,B). This suggests that a network receiving correlated input, for example from an upstream area, is more likely to generate chaotic dynamics than a network receiving static input without noise, which can potentially explain the prevalence of correlated activity across cortex.

**Figure 6:**
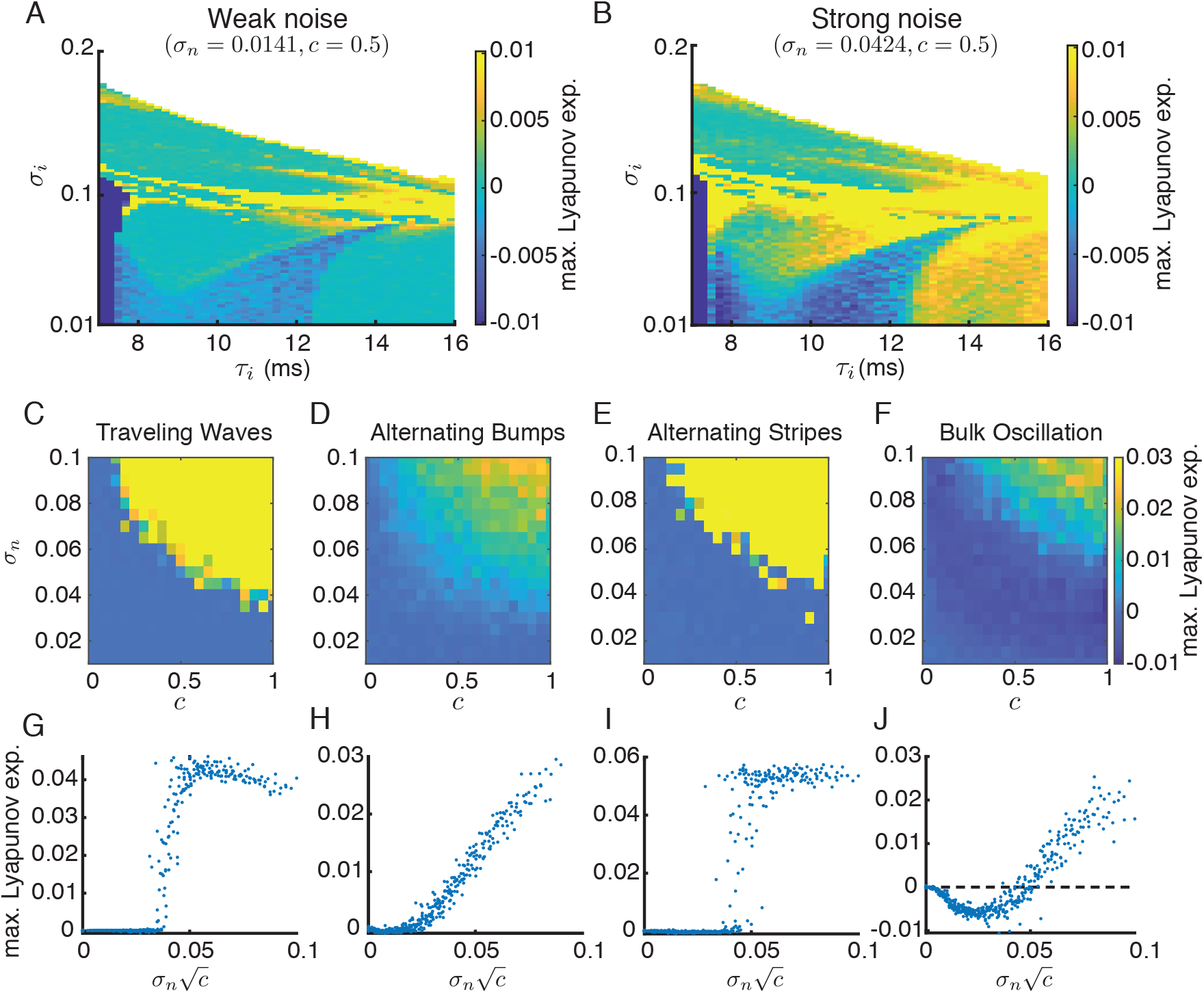
The maximal Lyapunov exponent (MLE) for the two-dimensional networks with input noise. **A-B**. The MLE map with weak (**A**, *σ*_*n*_ = 0.0141) and strong (**B**, *σ*_*n*_ = 0.0424) noise. The correlation of the input noise is *c* = 0.5. **C-F**. MLE as a function of the amplitude (*σ*_*n*_) and the correlation (*c*) of input noise, for four example networks that generate traveling waves (C; *τ*_*i*_ = 8, *σ*_*i*_ = 0.1, same as Fig. 3A), or alternating bumps (D; *τ*_*i*_ = 9, *σ*_*i*_ = 0.06, same as Fig. 3B), or alternating stripes (E; *τ*_*i*_ = 9, *σ*_*i*_ = 0.1, same as Fig. 3C) or bulk oscillation (F; *τ*_*i*_ = 8, *σ*_*i*_ = 0.03) without noise. **G-J**. Same as panels C-F with MLE as a function of the amplitude of the correlated noise component, 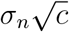.

In addition to inducing chaos, we find that noise can also synchronize bulk oscillations, reflected as negative MLE’s in the previously identified regime of bulk oscillations (light blue area in Fig. 6A-B). A negative MLE of a noise driven system means that networks starting at different initial conditions would converge to the same network dynamical trajectory that depends only on the noise realization (Baxendale, 1992; Jan, 1987).

Further, we investigate the impacts of the amplitude (*σ*_*n*_) and the correlation (*c*) of input noise on network dynamics. We compute the MLE of four examples of periodic solutions with varying *σ*_*n*_ and *c* (Fig. 6C-F). When there is no input noise (*σ*_*n*_ = 0), the MLE’s of these networks are zero, since they produce temporally periodic patterns. When driven by independent noise (*c* = 0), the MLE’s remain close to zero, which suggests that the periodic solutions are insensitive to independent noise. As *σ*_*n*_ and *c* increase, the MLE’s generally increase and become positive, indicating a transition to chaos (Fig. 6C-E). The bulk oscillation solution becomes synchronized with negative MLE for intermediate values of *σ*_*n*_ and *c* before transitioning to chaos (Fig. 6F).

We next measure how MLE depends on the amplitude of the correlated noise component, 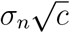 (Fig. 5A). When plotting as a function of 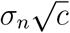, the MLE’s for varying *σ*_*n*_ and *c* collapse to a single function curve (Fig. 6G-J). This suggests that the MLE of the noise driven system mainly depends on the amplitude of the correlated noise component. The transition to chaos can be sharp, as in the cases of traveling waves and alternating stripes (Fig. 6G,I), or gradual, as in the cases of alternating bumps and bulk oscillation (Fig. 6D,F). Therefore, we identify that it is the correlated noise component that is responsible for inducing chaos and synchronizing bulk oscillations in the two-dimensional networks. In contrast, the network dynamical patterns are insensitive to independent noise.

## Discussion

Variability in neural responses is prevalent in cortex. The structure of the variability shared among a neuron population has important consequences on the information processing of the network (Kohn et al., 2016). However, the circuit mechanism underlying neural variability remains unclear. In this work, we discover a new dynamical regime in spatially distributed neuronal networks where spatiotemporal chaos produce large magnitude of shared variability in population activity. The statistical properties of the spatiotemporal chaos are consistent with population recordings from cortex, such as the broadband frequency power in single neuron responses, distance-dependent correlations and the low dimensionality of population responses.

Rate chaos in neuronal networks has been widely studied using random recurrent networks, where the connection weights from each neuron follow a Gaussian distribution with zero mean (Sompolinsky et al., 1988). The network solution transitions from a stable fixed point to chaos when the variance of connection weights exceeds a critical value. Similar transition to chaos is also observed in networks with separate excitatory and inhibitory populations (Kadmon and Sompolinsky, 2015) and in spiking neuron networks (Ostojic, 2014). In contrast, in spatially distributed networks, chaos appears after several solutions of different spatiotemporal patterns lose stability. As *σ*_*i*_ increases, the bulk oscillation solutions transition to alternating bump solutions, which transition to alternating stripes or traveling waves, and then to chaos (Fig. S1). Further theoretical analysis is needed to elucidate the path to chaos in spatial networks.

The spatiotemporal chaos in our networks has several distinct features from the chaotic solutions in random neuronal networks (Rajan et al., 2010; Schuecker et al., 2018; Sompolinsky et al., 1988). First, in networks of unstructured random connectivity, the correlations among neurons vanish as network size becomes large (scales as 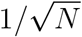 with *N* being the network size) (Rajan et al., 2010; Renart et al., 2010). Hence, those networks do not generate correlated population patterns, while the chaos in spatial networks produce distance-dependent correlations. Second, the dimensionality of population activity in random networks has been found to be high and increases linearly with network size (Engelken et al., 2020). This is in contrast with the low dimensional structure of the spatiotemporal chaos found in spatial networks (Fig. 4F) and population activity found in experimental data (Huang et al., 2019; Lin et al., 2015; Ruff et al., 2020; Schölvinck et al., 2015; Williamson et al., 2016). Lastly, previous work showed that time varying inputs, such as independent white noise and oscillatory inputs, suppress chaos (Engelken et al., 2020; Lajoie et al., 2013; Molgedey et al., 1992; Rajan et al., 2010; Schuecker et al., 2018), while we found that the chaotic solutions in spatial networks are insensitive to independent noise and that chaos can be induced by correlated noise (Fig. 6).

Several recent models have studied chaos in random neuronal networks with structured connectivity. For example, networks with a low rank connectivity component in addition to a random component can generate low dimensional coherent chaos, which can be utilized for complex computations (Landau and Sompolinsky, 2018; Mastrogiuseppe and Ostojic, 2018). Networks with cell-type-dependent distributions of connections can produce chaos with multiple modes of autocorrelation functions of individual neurons (Aljadeff et al., 2015). In this work, we demonstrate that networks with two-dimensional spatial couplings and no random connectivity can also generate chaos which resides in a low-dimensional state space. How random connectivity in combination with spatially ordered connectivity affect chaos remain to be studied in future work.

Chaotic dynamics in neuronal networks offer a rich “reservoir” of population activity patterns, which can be utilized to learn a target output function or accomplish complex neural computations (Buonomano and Maass, 2009; Lukoševičius and Jaeger, 2009; Pyle and Rosenbaum, 2017; Schuessler et al., 2020; Sussillo and Abbott, 2009). Near the transition of chaos, networks can generate slow dynamics which are important for temporal integration and necessary for many behavioral tasks (Huang and Doiron, 2017; Toyoizumi and Abbott, 2011). Here we find a new type of spatiotemporal chaos in networks where the connectivity features are consistent with cortical anatomy. It would be fruitful to explore the computational benefits of such chaos in spatially distributed networks.

## Methods

### Stability of fixed point solutions

Linearization around the fixed point solution 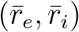 of Eqs 1-2 in Fourier space gives a Jacobian matrix at each spatial Fourier mode:

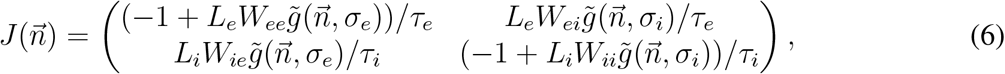

Where 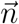 is the Fourier mode, 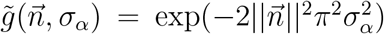 is the Fourier Gaussian kernel, and 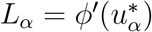 evaluated at the fixed point 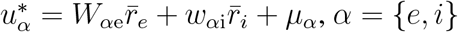.

The fixed point is stable if all eigenvalues of 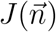 have negative real part for any 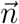 (Fig. 2B). Note that the stability only depends on the wave number 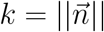.

### Stability of bulk oscillation solutions

To analyze the stability of bulk oscillation solutions, we linearize around the time dependent limit cycle solution, 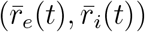, and obtain the Jacobian matrix at each Fourier mode, which is also periodic in time (Ali et al., 2016; Teschl, 2012):

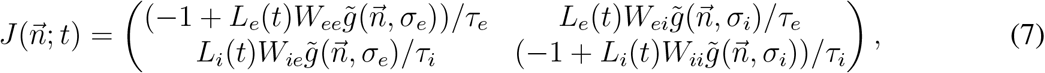

where 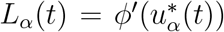 evaluated along the limit cycle solution 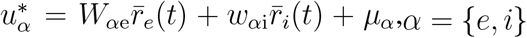. The perturbation of rate, 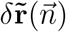, at Fourier mode, 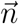, follows a linear system with periodic coefficients:

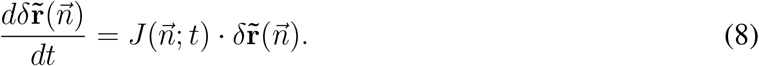

We obtain a principal fundamental matrix solution of Eq. 8 by solving 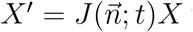 with initial conditions *X*(0) = *I*, where *I* is the identity matrix. The stability of the bulk oscillation solution is then determined by the eigenvalues of the Monodromy matrix,

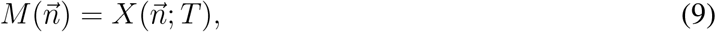

where *T* is the period of the bulk oscillation solution. If any of the eigenvalues of 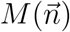 have magnitude greater than 1 at some Fourier model 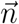, then the bulk oscillation will lose stability with spatial mode 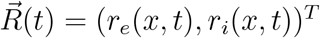. A real eigenvalue less than −1 indicates a period-doubling bifurcation, while a real eigenvalue larger than 1 suggests a pattern formation with the same period as the bulk oscillation (Ali et al., 2016).

### Maximal Lyapunov exponent

The maximal Lyapunov exponent (MLE) is computed numerically using the method by Wolf et al. (1985). We first simulate the network long enough such that the solution has converged to an attractor. We denote the solution trajectory as 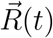.

We continue simulating the trajectory for *n* time points, {*t*_0_, *t*_1_, …, *t*_*n*_}, with step size Δ*t*. At the initial time point, *t*_0_, we perturb the trajectory, 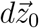, by 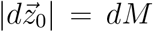, which is in a random direction and has a small magnitude 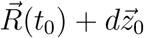. We integrate the same model system with the perturbed initial condition, 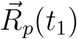, by one time step, Δ*t*, and obtain the perturbed trajectory, 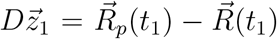. The separation between the two trajectories is 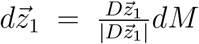. Then we choose the perturbation at the next time step as 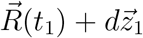. In this way, the perturbation in each time point has a magnitude of *dM*, and is in the same direction as the separation between the original and the perturbed trajectories at the previous time step. We then integrate the same model system with the perturbed initial condition, 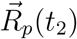, by one time step and obtain 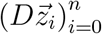. We repeat this procedural *n* steps and obtain a sequence of trajectory separations, 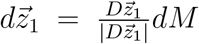. Lastly, the Maximal Lyapunov exponent is computed as

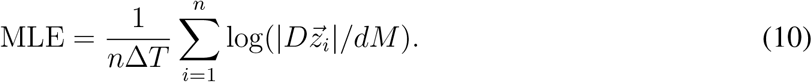

### Spatial correlation

The spatial correlations in Fig. 4B-C are defined as the Pearson correlation coefficient of *r*_*e*_, for each neural distance (Δ*x*, Δ*y*):

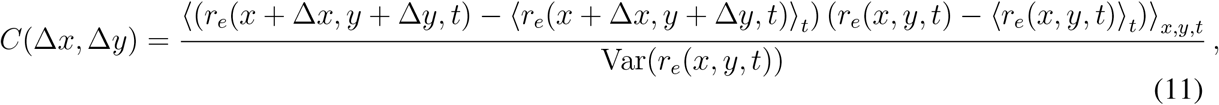

where ⟨·⟩_*t*_ is average over time and ⟨· ⟩_*x,y,t*_ denotes average over time and space.

### Power spectrum

The power spectrum in Fig. 4D is defined as:

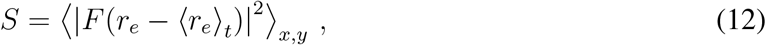

where the function *F* is the Fourier transform in time.

### Input noise

The input noise terms 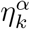 and 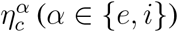 in Eq. 5 are modeled as independent Ornstein–Uhlenbeck processes:

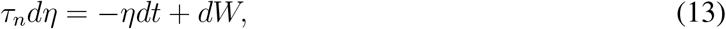

where *W* is a Wiener process, and the noise time constant is *τ*_*n*_ = 5 ms.

### Network parameters

Unless specified otherwise, the network parameters are *W*_*ee*_ = 80, *W*_*ei*_ = −160, *W*_*ie*_ = 80, *W*_*ii*_ = −150, *τ*_*e*_ = 5 ms, *σ*_*e*_ = 0.1, *µ*_*e*_ = 0.48 and *µ*_*i*_ = 0.32. *τ*_*i*_ and *σ*_*i*_ vary in each figure and are specified in figure axes and captions. The number of neurons in the two dimensional network are *N* = 100 × 100 for each of the excitatory and inhibitory populations. In the one dimensional network *N* = 100 for each of the populations.

## Acknowledgments

C.H. was supported by NIH grant R01NS121913 and the University of Pittsburgh. B.E. was partially supported by NSF DMS 1951099. We thank Daniele Avitabile for his code and discussion regarding computing the stability of the traveling wave regime (Fig. 2B). We thank Jonathan Kadmon for the consultation regarding the MLE computation.

## Author Contributions

N.M., B.E. and C.H. conceived the project; N.M. performed the simulations and analysis; C.H. supervised the project; all authors contributed to writing the manuscript.

## Data and Software Availability

Computer code for all simulations and data analysis is available upon request.

## Supplemental Information

**Figure S1:**
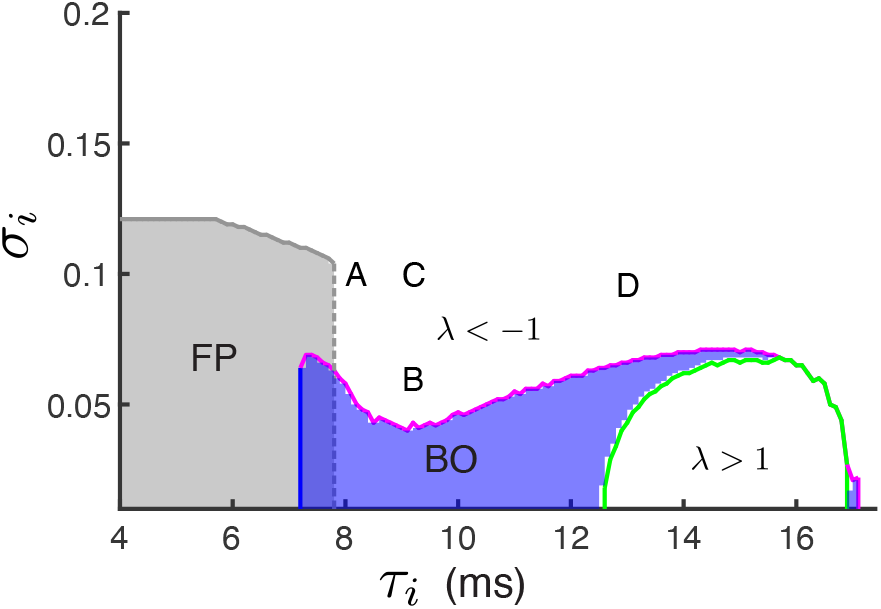
Related to Figure 3. Phase diagram of networks with two dimensional spatial coupling. Same format as Fig. 2B in the main text. The letters mark the locations of the parameters used in Fig. 3A-D.

**Figure S2:**
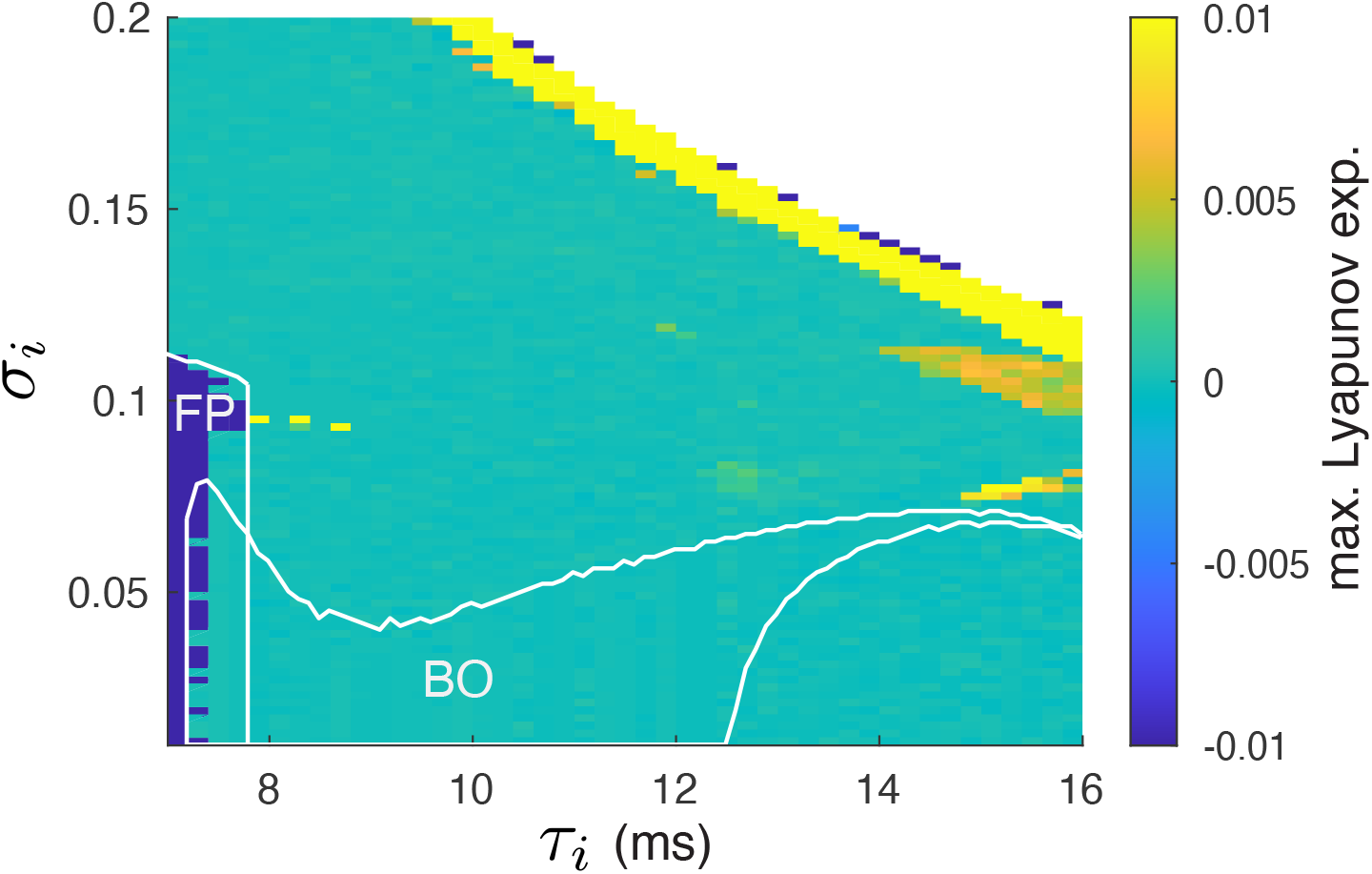
Related to Figure 4. The maximal Lyapunov exponents of networks with one dimensional spatial coupling. Same format as Fig. 4A. The maximal Lyapunov exponent as a function of the projection width (*σ*_*i*_) and the time constant (*τ*_*i*_) of the inhibitory neurons. The white curves are the stability borders of the parameter regions of stable fixed point (FP) and bulk oscillation (BO) solutions (same regions as in Fig. 2B).

